# Carbonate bedrock may not alleviate DIC limitation of snow algae — a test of hypothesis in the Medicine Bow Mountains, WY, USA

**DOI:** 10.1101/2023.02.24.529969

**Authors:** Trinity L. Hamilton, Jeff Havig

## Abstract

Snow is a critical component of the Earth system. High elevation snow can persist into the melt season and hosts a diverse array of life including snow algae. Due in part to the presence of pigments, snow algae lower albedo and accelerate snow melt which has led to increasing interest in identifying and quantifying the environmental factors that constrain their distribution. Dissolved inorganic carbon (DIC) concentration is low in supraglacial snow on Cascade stratovolcanoes and snow algae primary productivity can be stimulated through DIC addition. Here we asked if CO_2_ would still be a limiting nutrient for snow hosted on glacially eroded carbonate bedrock (which could provide an additional source of DIC). We assayed snow algae communities for nutrient and DIC limitation on two seasonal snowfields on glacially eroded carbonate bedrock in the Snowy Range of the Medicine Bow Mountains, Wyoming, USA. DIC stimulated snow algae primary productivity in snow with lower DIC concentration despite the presence of carbonate bedrock, which alleviated DIC limitation in the other site. Our results support the hypothesis that increased atmospheric CO_2_ concentrations may lead to larger and more robust snow algae blooms globally, even for sites with carbonate bedrock.

## INTRODUCTION

Snow is an important part of earth’s biogeochemical cycles (Anesio et al., 2017). When snowpack persists into the melt season (in late spring, summer, and early fall), diverse assemblages of microbial life including archaea, bacteria, and microbial eukaryotes can persist on snow surfaces due to the presence of liquid water. The most recognizable microbial life on snow are snow algae — eukaryotic photosynthetic algae typically within the within the Chlorophyceae. Snow algae are common primary producers on snowpack where they fix inorganic carbon into biomass using sunlight. As phototrophs, they make pigments and carotenoids to harvest sunlight. Snow algae also commonly make astaxanthin, an intracellular carotenoid that likely serves as an antioxidant under high UV and visible light conditions protecting the algae’s photosynthetic apparatus from damage. Astaxanthin colors the snow algae cells red, and thus snow algae blooms are commonly observed as red or watermelon colored snow (Figure 1). In snow algae, the absorption spectra of the red astaxanthin carotenoids results in a reduction of the albedo of snow they inhabit, accelerating melt and increasing the amount of liquid water available for biological processes (for a recent review see Hotaling et al., 2021). Because snow algae are the most common primary producer on snow and their presence accelerates melt — red snow algae decreased albedo in the Arctic by as much as 13% (Lutz et al., 2016) and increased snowmelt by ~21% on an Alaskan icefield (Ganey et al., 2017) — understanding the biotic and abiotic factors that control their distribution is of keen interest.

**Figure 1.**
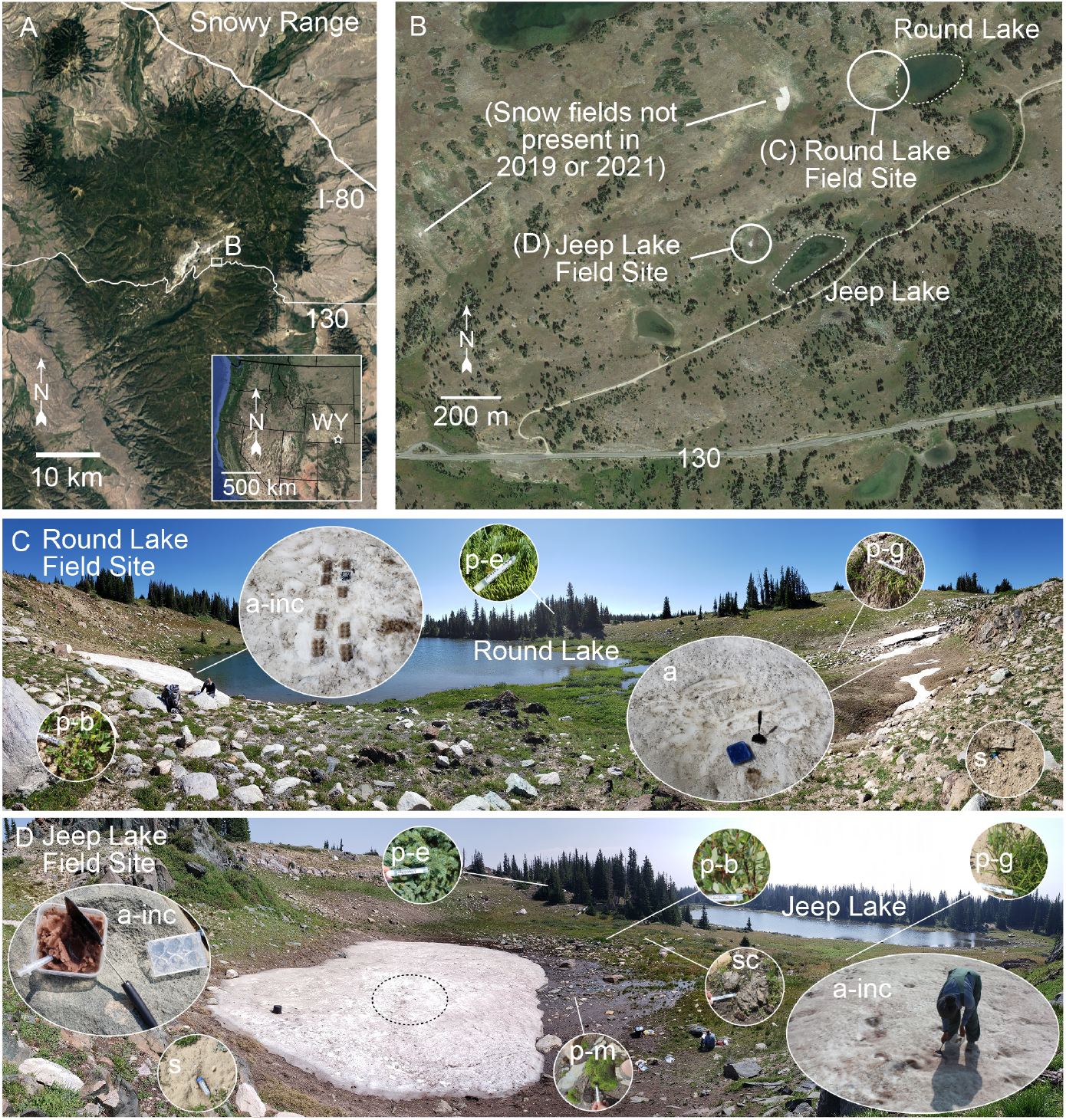
Sampling location. A) Satellite image of the Snowy Mountain Range in the Medicine Bow Mountains, WY. Inset shows the western United States, with an open star indicating the location of the Snowy Mountain Range. Open rectangle indicates the area shown in panel B. B) Zoomed in satellite image of the sampling area indicating the locations near Round Lake (with images of the sample site shown in panel C) and Jeep Lake (with images of the sample site shown in panel D). Snow fields that were visible in the satellite image but were not present at the time of sampling are noted. C) Panoramic image of the Round Lake field site looking east with the two snow algae sample sites indicated. Inset images show snow algae samples being collected for carbon uptake experiments (a-inc) or only geochemistry and molecular samples (a), as well as examples of contextual samples including plants (evergreen trees (p-e), broadleaf plants (p-b), and grasses and sedges (p-g)) and soil (s). D) Panoramic image of the Jeep Lake field site looking east with the snow algae incubation site indicated by a dashed open black circle. Inset images show snow algae samples being collected for carbon uptake experiments (a-inc), as well as examples of contextual samples including plants (evergreen trees (p-e); broadleaf plants (p-b), moss (p-m), and grasses and sedges (p-g), soil, (s), and scat (sc).

In general, nutrients are thought to limit primary productivity on snow and glacier ice (Wolff, 2013; Williams et al., 2016; McCutcheon et al., 2021) and microbial activity on snow and ice relies on the delivery of exogenous nitrogen, phosphorus, and trace elements (Lutz et al., 2015; Hamilton and Havig, 2017; Havig and Hamilton, 2019). In the Cascade Range of the US, wind-blown volcanic ash and rock flour are important sources of trace metals and phosphorous for snow algae communities while precipitation is an important source of fixed nitrogen (Hamilton and Havig, 2017). Location-specific nutrient delivery has been observed elsewhere. For example, recently erupting volcanoes in Iceland deliver fresh ash to nearby glaciers. The fresh volcanic ash is an important source of essential nutrients (Lutz et al., 2015). Studies of supraglacial snow on stratovolcanoes in the Cascade Range of the Pacific Northwest or glaciers near erupting volcanos may lead to unique observations compared to glaciers that override granitic or carbonate-rich bedrock.

Due to human activity, atmospheric CO_2_ was above 417 ppm as of 2022 (Friedlingstein et al., 2022). Coincident with increasing CO_2_, average global temperatures have increased by over 1°C as of 2021 (relative to the 1850-1900 averaged value), with best estimates of an additional 0.5 °C increase by 2040, driving recent and future glacial retreat and receding snowpack (Arias et al., 2021). Alpine ecosystems are particularly susceptible to a changing climate and in the Rocky Mountain Range, mountain glaciers are rapidly retreating. Climate change will likely alter precipitation regimes (changing seasonal snowfall and rain (e.g., East and Sankey 2020)) and impact the delivery of material to snow and ice surfaces. Land-use change, urbanization, and changes in agricultural nitrogen deposition have increased levels of fixed nitrogen in high elevation ecosystems across the Rocky Mountain Range (Brahney et al., 2015) while phosphorus, heavy metals, and carbonates are also delivered via dry fall of dust from the erosion of desert and agricultural soils (Lawrence and Neff, 2009). Forest fires also increase nutrient deposition including nitrogen and phosphorus (Bladon et al., 2014).

Despite the apparent lack (or low concentration) of fixed nitrogen, phosphorous, and essential trace elements in snow, snow algae productivity is not always stimulated by the addition of these nutrients. In our previous studies of snow algae on supraglacial snow on stratovolcanoes in the Cascade Region of the US, we observed an ambiguous response of snow algae to fixed nitrogen (in the form of NO_3_- or NH_4_+) or phosphate — these nutrients did not uniformly stimulate photosynthesis in snow algae from three different stratovolcanoes (Hamilton and Havig, 2017). However, snow, ice, and proglacial streams were undersaturated with respect to dissolved CO_2_ leading us to hypothesize the snow algae were limited in inorganic carbon. Indeed, the addition of inorganic carbon stimulated photosynthesis (Hamilton and Havig, 2020).

The low availability of dissolved inorganic carbon (DIC) in supraglacial snow environments on stratovolcanoes are due in part to the local bedrock – silicic volcanic rocks are effectively devoid of inorganic carbon, constraining CO_2_ availability to either allochthonous atmospheric input or release from autochthonous heterotrophic breakdown of organic matter as primary sources of inorganic carbon for photoautotrophy (Havig and Hamilton, 2019). What remains poorly constrained is the impact of carbonate-rich bedrock dissolution increasing DIC availability (e.g., CO_2_ + H_2_O + CaCO_3_ => 2 HCO_3_- + Ca^2+^) on snow algae productivity. To test this, we replicated the experiments conducted on supraglacial snow algae communities on Cascade stratovolcanoes on snowfield environments in the Snowy Range of the Medicine Bow Mountains in Wyoming, USA – a location with glacially eroded carbonate bedrock that possess a uniquely positive δ^13^C signal – to test for DIC vs. nutrient (P, fixed N) stimulation of snow algae productivity. We examined microbial community composition and the response of photosynthesis to the addition of essential nutrients including fixed nitrogen, phosphate, iron, and dissolved inorganic carbon in two snowfields in the Snowy Range overlying the Nash Creek Formation carbonate.

## MATERIALS AND METHODS

### Sample Site Description

The Snowy Mountain range (or the Heneecee Niicie Hoheni to the Arapaho) is located in south-central Wyoming and consists of Precambrian (~ 2.9 to 1.7 Ga) metasedimentary and metavolcanic units thrust onto Phanerozoic rocks during the ~80 to 55 Ma Laramide orogeny (Houston and Karlstrom, 1992). The elevations of the Snowy Mountain range from the base of ~2500 m (~8100 ft) to the highest point at the apex of Medicine Bow Peak at 3663 m (12,018 ft). The sample sites selected were situated on the ~ 2.0 Ga Nash Fork Formation carbonate with an elevation of ~3250 m (10,560 ft). The Nash Fork Formation records a period of extended positive isotope excursion following the Great Oxidation Event (~2.45 Ga), with the section underlying the sampling areas having carbonates with δ^13^C values of 6.7 (± 1.2) ‰ (Bekker et al., 2003). The vegetation is predominantly alpine vascular plants (e.g., grasses, sedges, and broadleaf plants) as well as local patches of non-vascular plants (mosses). Copses of spruce (*Picea engelmannii*) and fir (*Abies lasiocarpa*) were located around the sampling areas, as well as thickets of willow (*Salix* spp.). Most precipitation received by the Snowy Range is in the form of snow from October-May: 50-80% of annual precipitation which can be distributed heterogeneously by strong west wind (Hiemstra et al., 2004), topography, and vegetation. Thus, seasonal snow persists well into the summer months at high elevations in the range and snow algae are commonly observed on these snow patches. In addition to the presence of sedimentary rocks with carbonates for bedrock at the field sites in the Snowy Range, there has been increased P and fixed N deposition observed across the Rocky Mountains (Brahney et al., 2015). Seasonal snowfields in the Snowy Range are also distinct from supraglacial snow because they blanket vegetated and (often) developed soils.

### Sample Collection, RNA Extraction, and Sequencing

Samples were collected from snowfields in the Medicine Bow Mountains in August 2019 and 2021. Snow for geochemical and biological analyses was collected by scraping the upper ~3 cm of snow into a clean 1 L polypropylene bottle (soaked in 10% trace element grade HNO3 for three days, triple rinsed with 18.2 MΩ/cm deionized water). Snow was thawed in a closed bottle with minimal atmospheric exposure. Melted snow or water (stream or lake) was filtered through 25 mm diameter 0.2 μm polyethersulfone syringe filters (VWR International, Radnor, PA) and distributed into bottles for geochemical analysis as described. Temperature and pH were measured in the melted snow using a WTW 330i meter and probe (Xylem Analytics, Weilheim, Germany). Samples for RNA extraction and stable isotope analyses were collected from the melted snow using sterile pipettes, transferred to sterile tubes and immediately frozen on liquid N_2_ and stored at −80°C until further processed.

Samples for induced coupled plasma optical emission spectroscopy (ICP-OES) and induced coupled plasma mass spectrometry (ICP-MS) were filtered into a 15 mL centrifuge tubes that had been acid washed (soaked in 10 % trace element clean HNO3 for three days, triple rinsed with 18.2 MΩ/cm deionized water), acidified (200 μL of concentrated OmniTrace Ultra™ concentrated nitric acid (EMD Millipore, Billerica, MA)), and kept on ice/refrigerated at 4 °C in the dark until analysis. ICP-OES analysis for Ca, Na, K, Mg, P, and Si, and ICP-MS analysis for Al, Ti, Mn, and Fe was conducted at the Quantitative Bio-element Imaging Center at Northwestern University (QBIC) at Northwestern University. For anions, water was filtered into 15-mL centrifuge tubes (pre-soaked in 18.2 MΩ/cm DI water). Ion chromatography analyses of anions was performed by the QBIC at Northwestern University.

Samples for dissolved inorganic carbon (DIC) analysis were filtered into a gas-tight syringe and then injected into Labco Exetainers® (Labco Limited, Lampeter, UK) pre-flushed with He, with excess He removed following introduction of 8 mL of filtered sample with minimal agitation. Samples were stored inverted until and on ice/refrigerated at 4 °C until returned to the lab, where 1 mL of concentrated H3PO4 was added and the samples shipped to the Stable Isotope Facility (SIF) at the University of California, Davis for analysis. DIC analysis for concentration and ^13^C isotopic signal using a GasBench II system interfaced to a Delta V Plus isotope ratio mass spectrometer (IR-MS) (Thermo Scientific, Bremen, Germany) with raw delta values converted to final using laboratory standards (lithium carbonate, δ^13^C = - 46.6 ‰ and a deep seawater, δ^13^C = +0.8 ‰) calibrated against standards NBS-19 and L-SVEC.

For dissolved organic carbon (DOC) analysis, 0.2 μm polyethersulfone syringe filters were preflushed with ~ 30 mL of sample. After, ~ 40 mL were collected into a 50 mL centrifuge tube and then immediately flash-frozen on dry ice and kept frozen and in the dark until analysis at the SIF. DOC analysis for concentration and ^13^C isotopic signal were carried out using O.I. Analytical Model 1030 TOC Analyzer (O.I. Analytical, College Station, TX) interfaced to a PDZ Europa 20-20 isotope ratio mass spectrometer (Sercon Ltd., Cheshire, UK) utilizing a GD-100 Gas Trap Interface (Graden Instruments) for concentration and isotope ratio determination with raw delta values converted to final using laboratory standards (KHP and cane sucrose) calibrated against USGS-40, USGS-41, and IAEA-600.

### CO_2_ photoassimilation and C and N natural abundance

Inorganic carbon uptake was assessed *in situ* using a microcosm-based approach through the addition of NaH^13^CO_3_. Snow was collected from the surface using a pre-sterilized spatula, placed into a clean container, and allowed to melt to a slush-slurry. An equal volume of snow slurry (~ 10 mL) or was then transferred into each well of a sterile six well tray and the lid placed over the tray. Assays were initiated by addition of NaH^13^CO_3_ (Cambridge Isotope Laboratories, Inc., Andover MA).

We assessed the potential for photoautotrophic (light) and chemoautotrophic (dark) NaH^13^CO_3_ uptake (100 μM final concentration). To assess CO_2_ assimilation in the light, six well trays (n = 1 per site) were amended with NaH^13^CO_3_ and incubated under full sun. To assess CO_2_ assimilation in the dark, six well trays (n = 1 per site) were amended with NaH^13^CO_3_ and completely wrapped in aluminum foil. To assess nutrient limitation, a series of microcosms were amended with fixed nitrogen (KNO_3_ (100 μM final concentration)) or NH_4_Cl (100 μM final concentration)), PO_4_^3-^ (as KH_2_PO_4_, 100 μM final concentration), or Fe (as FeSO4-7H2O, 500 nM final concentration following Zhang et al., 2019). In addition, we assessed CO_2_ limitation by performing the microcosms at 250 μM dissolved inorganic carbon (NaHCO_3_). *in situ* microcosms were performed under full sun between 11 am and 3 pm. Following incubation time (t ≈ 2.5 hours), incubation trays were flash frozen on dry ice and kept frozen until processed.

Following return to the lab, microcosm and ^13^C natural abundance samples were rinsed with 1 M HCl to remove any carbonate minerals and any residual NaH^13^CO_3_ label, triple rinsed with 18.2 MΩ/cm deionized water and dried (8 h, 60 °C). ^15^N samples were dried (but not acidified). Dried samples were ground and homogenized using a clean mortar and pestle (silica sand and 80 % ethanol slurry ground between samples, then mortar and pestle rinsed with ethanol followed by 18.2 MΩ/cm deionized water and dried). Ground samples were weighed and placed into tin boats, sealed, and analyzed via a Costech Instruments Elemental Analyzer (EA) periphery connected to a Thermo Scientific Delta V Advantage Isotope Ratio Mass Spectrometer (IR-MS) by the SIF at UC Davis. Reported values of DIC uptake (carbon fixation rates) reflect the difference in uptake between the labeled assays and unlabeled biomass.

To characterize the C and N isotopic values of snow algae microbial communities and differentiate between allochthonous and autochthonous organic material, contextual samples from on and near the snowfield were collected including plants, plant material on the snow, and surrounding sediments. Contextual samples were treated and analyzed as described above.

### RNA extraction and sequencing

Total RNA was extracted from samples using a RNeasy PowerSoil Total RNA Kit. Following purification, RNA was subjected to DNAse I digestion (Roche, Indianapolis, IN, USA) for 1 h at room temperature (~22 °C) and then further purified using a High Pure RNA Isolation Kit (Roche, Basel, Switzerland) and was stored at −80 °C in a solution of 100% ethanol and 0.3 M sodium acetate until further processed. The concentration of RNA was determined using a Qubit RNA Assay kit (Molecular Probes, Eugene, OR, USA) and a Qubit 3.0 Fluorometer (Life Technologies, Carlsbad, CA). RNA extracts were screened for the presence of contaminating genomic DNA by performing a PCR using ~1 ng of RNA as template and the same primers used for sequencing (described below). RNA was extracted from duplicate samples and DNA-free RNA was pooled for each sample. cDNA was synthesized from 20 ng of purified RNA using the Superscript IV Reverse Transcriptase (Thermo Fisher Scientific, Waltham, MA) according to the manufacturer’s instructions and purified by ethanol precipitation. cDNA was re-suspended in nuclease-free water for use in amplicon sequencing. Total cDNA was submitted to the University of Minnesota Genomics Center (UMGC) where libraries dual indexed Nextera XT DNA were prepared following Gohl et al 2016. Amplicons were sequenced at UMGC using MiSeq Illumina 2 x 300 bp chemistry with the primers 515Ff and 806rB targeting the V4 region of bacterial and archaeal 16S SSU rRNA gene sequences (Caporaso et al., 2012; Apprill et al., 2015) and primers E572F and E1009 targeting the V9 region of eukaryotic 18S SSU rRNA gene sequences (Comeau et al., 2011).

### Sequence analysis

Post sequence processing was performed using the mothur (ver. 1.44.0) sequence analysis platform (Schloss et al., 2009) following the MiSeq SOP (Kozich et al., 2013) as described previously (e.g. Havig and Hamilton, 2019). Briefly, read pairs were assembled and resulting contigs with ambiguous bases were removed and trimmed to include only the overlapping regions. Chimeras were identified and removed using UCHIME (Edgar et al., 2011) from all the data sets. 16S rRNA amplicons were aligned against the SILVA v138 database and clustered into operational taxonomic units (OTUs) at a sequence similarity of 0.97. 16S rRNA OTUs were classified in mothur using the SILVA database (v138). 18S rRNA amplicons were aligned against the PR^2^ database (ver. 4.12.0) (Guillou et al., 2013). 18S rRNA OTUs were clustered into OTUs at a sequence similarity of 0.99 and classified in mothur using the PR^2^ database. All post-mothur processing was carried out in R (ver. 4.0.3; R Core Team, 2018)

## RESULTS AND DISCUSSION

Snow is typically low in aqueous nutrients including fixed nitrogen, phosphorus, and iron (Wolff, 2013; Williams et al., 2016; Havig and Hamilton 2019; McCutcheon et al., 2021). In previous studies on supraglacial snow on stratovolcanoes in the Pacific Northwest region of the US, we tested the hypothesis that snow algae communities are nutrient limited. PNW supraglacial snow is typically above the treeline and the surrounding landscape is predominantly recently deglaciated sediments sparsely populated by mosses and pioneering species of vascular plants. Our data indicated supraglacial snow algae in the PNW were not nutrient limited and instead these communities assimilate fixed nitrogen from precipitation and phosphorus and iron from local bedrock (Hamilton and Havig 2017; Havig and Hamilton 2019). Here, we expanded this study to examine nutrient limitation on seasonal snow in the Snowy Range of the Medicine Bow Mountains, WY, USA. These snow fields are distinct from PNW sites — they override developed soils and are surrounded by trees, shrubs, grasses, and other foliage, and the bedrock is the Nash Creek Formation (a ~ 2.0 Ga marine-deposited carbonate)(Houston and Karlstrom, 1992). Late season snow was collected from seasonal snowfields in the Snowy Range of the Medicine Bow Mountains, WY, USA near Round Lake in August of 2019 and Jeep Lake in August of 2021 (Figure 1). We targeted areas of snow that were visibly pigmented (orange snow near Round Lake and pink/red snow near Jeep Lake). Additional samples from the proximal meltwater streams and the lakes were also collected for comparison.

### Snow algae

Snow algae composition was distinct between Round Lake and Jeep Lake snowfields despite the lakes being within ~500 m of each other (Figure 1). OTUs affiliated with the *Chlorococcales* were abundant in Round Lake (yellow to orange) snow while the most abundant snow algae OTUs in pink snow from Jeep Lake included members of the *Chlamydomonadales* and *Chloromonas* (Figure 2). Small numbers of *Chlorococcales* were also present in Jeep Lake snow. Algae composition was also distinct from lake samples: OTUs affiliated with Ochrophytina *Nannochloropsis oceanica* and the diatom *Navicula cryptotenlla* were abundant in lake sediments. We also sampled meltwater streams directly downstream from both snow patches. At Round Lake, 18S rRNA OTUs were largely distinct between snow, stream, and lake. In the meltwater stream, OTUs affiliated with *Chloromonas pichinchae* were abundant while these OTUs comprised a much small fraction of the 18S rRNA community in snow and lake samples. In contrast, *Chloromonas* OTUs were abundant in both Jeep Lake snow and the meltwater stream. Meltwater streams could act as conduits of dispersal for snow algae. This observation is supported at Jeep Lake (based on similarity between snow and stream 18S rRNA communities) but not at Round Lake; however, we sampled sediments and not meltwater thus the conditions in the sediments and other characteristics including stream flow rates likely control the composition of sediment algal communities. Indeed, algal composition was distinct between lake sediments and biofilms and snow.

**Figure 2.**
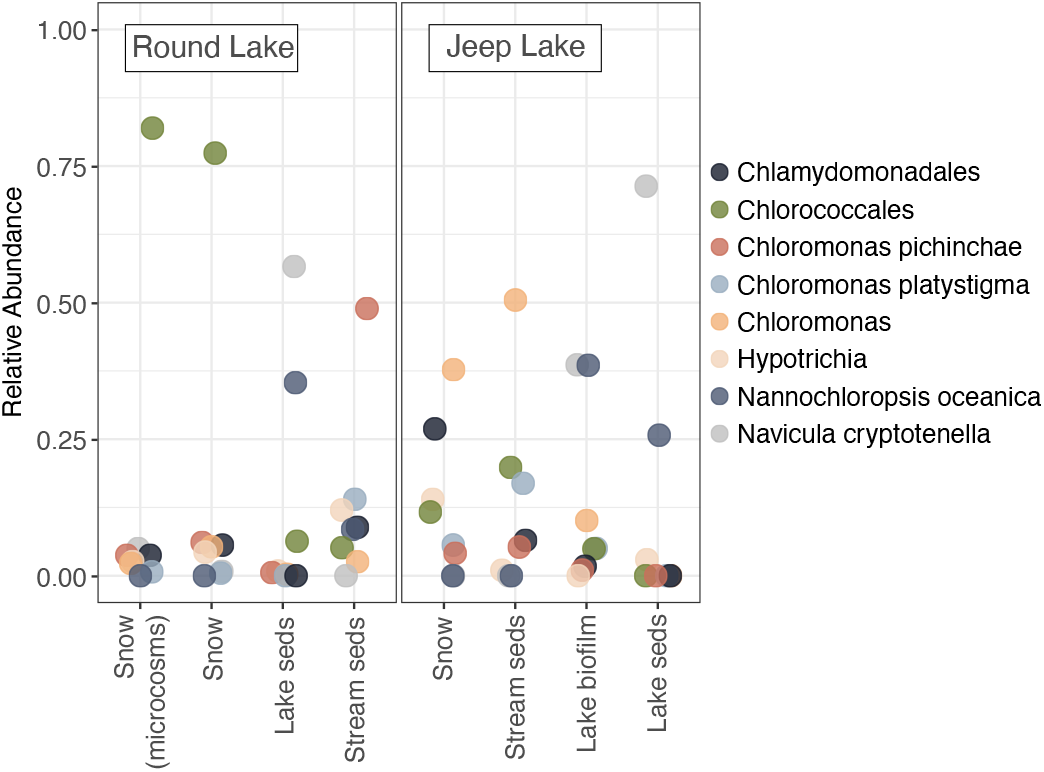
Relative abundance of algal small subunit rRNA (18S rRNA) OTU transcripts recovered from snow, lake and stream sediments (seds) and lake biofilms. OTUs for each library were binned at the genus level. The Round Lake sample that corresponds to the site where microcosms were performed is indicated.

Other biotic and abiotic factors might impact snow algae composition. For example, the Round Lake snow (from 2019) had a pH of 6.0 to 6.2 while Jeep Lake (from 2021) snow had a pH of 7.8 (Table 1). The difference in pH values for the Round Lake versus Jeep Lake snow algae sample sites are driven by inorganic carbon sources. Round Lake snow likely had minimal input of carbonate-bearing dust present on the snow surface at the time of sampling, making input of atmospheric CO_2_ the primary source of dissolved inorganic carbon (DIC) (e.g., CO_2_(g) + H_2_O => H_2_CO_3_ => H^+^ + HCO_3_-). This leads to the relatively low DIC concentration (≤ 41 μmol/L C) and mildly acidic pH in the Round Lake snow samples. These values are consistent with those reported for PNW snow and snow runoff samples, which had average an DIC concentration of 23.5 (±11.5) μmol/L C and an average pH value of 5.4 (± 0.8) (Hamilton and Havig, 2018; Havig and Hamilton, 2019). Conversely, the Jeep Lake snow algae site had a DIC value of 131 μmol/L C, and with a pH value of 7.8 was buffered by carbonate mineral dissolution (e.g., CO_2_(g) + H_2_O + CaCO_3_ => Ca^2+^ + 2 HCO_3_-).

**Table 1.** GPS coordinates, pH, conductivity, temperature, aqueous geochemistry, and stable isotope analyses. Dissolved inorganic carbon (DIC); dissolved organic carbon (DOC). Values in grey are near the limits of calibration curves, and should be taken as approximations.

| Sample Type | GPS |  | Elev.<br>(m) | pH | DIC |  | DOC |  |
| --- | --- | --- | --- | --- | --- | --- | --- | --- |
| | 13 T<br>E | UTM<br>N | | | $\mu\text{M}$ | $\delta^{13}\text{C}$<br>‰ | $\mu\text{M}$ | $\delta^{13}\text{C}$<br>‰ |
| Round Lake |  |  |  |  |  |  |  |  |
| Snow algae | 393740 | 4579422 | 3240 | 6.2 | 16.30 | -14.78 | 194.77 | -25.77 |
| Snow algae | 393686 | 4579354 | 3242 | 6.0 | 40.84 | -19.83 | 278.69 | -25.53 |
| Lake | 393747 | 4579412 | 3236 | 8.1 | 1627.15 | -7.94 | 186.17 | -25.10 |
| Stream | nd | nd | nd | 8.5 | 99.92 | -10.31 | 83.21 | -25.63 |
| Jeep Lake |  |  |  |  |  |  |  |  |
| Snow algae | 393395 | 4579012 | 3285 | 7.8 | 130.69 | -24.01 | 302.32 | 302.32 |
| Stream | nd | nd | nd | 7.5 | 179.40 | -11.00 | 27.72 | 27.72 |
| Lake | 393466 | 4578952 | 3246 | 8.1 | 616.62 | -11.11 | 363.43 | 363.43 |
| | SO<sub>4</sub><sup>2-</sup><br>$\mu\text{M}$ | P<br>$\mu\text{M}$ | Ca<br>$\mu\text{M}$ | Mg<br>$\mu\text{M}$ | Mn<br>nM | Fe<br>nM | | |
| Round Lake |  |  |  |  |  |  |  |  |
| Snow algae | 0.67 | 0.54 | 1.81 | 2.55 | 87.82 | 1326.27 |  |  |
| Snow algae | bdl | 0.46 | 4.03 | 1.23 | 112.03 | 474.36 |  |  |
| Lake | 5.36 | 0.14 | 459.42 | 377.43 | 65.90 | 1010.68 |  |  |
| Stream | 1.17 | 0.34 | 5.53 | 3.09 | 70.11 | 107.52 |  |  |
| Jeep Lake |  |  |  |  |  |  |  |  |
| Snow algae | 3.01 | 3.30 | 10.06 | 2.30 | 467.48 | 390.15 |  |  |
| Stream | 2.98 | 4.96 | 40.57 | 8.93 | 62.60 | 175.63 |  |  |
| Lake | bdl | bdl | 302.35 | 163.83 | 8.19 | 82.63 |  |  |

DIC in both sites is likely sourced from a combination of sources, including atmospheric CO_2_(g), dissolution of Nash Fork Formation carbonate, and heterotrophic breakdown of organic carbon (C_org_). For this combination of sources, we would predict a range of DIC δ^13^C values from −15.5 ‰ (assuming DOC as the source of CO_2_) to −17.4 ‰ (assuming soil as the source of CO_2_) to −19.2 ‰ (assuming plant matter as the source of CO_2_). DIC concentration and δ^13^C values along with pH are consistent with Round Lake snow fields having atmospheric CO_2_ as a primary source for DIC (low DIC concentration, slightly acidic pH) with minimal carbonate dissolution while Jeep Lake snow fields had an alkaline pH value and higher DIC concentration due to dissolution of locally-derived carbonate minerals. These conclusions are based on the following evidence:

1. Assuming an atmospheric δ^13^C signal of −8 ‰ for CO_2(g)_ and DIC at equilibrium at a temperature of ~ 0 °C, we would predict a DIC δ^13^C value of −9.2 ‰ (Mook et al., 1974; Havig and Hamilton, 2019) which is heavier than our snow samples.
2. The Nash Fork Formation carbonate forms the bedrock at the Round and Jeep Lake sites (Houston and Karlstrom, 1992), and the section of the carbonates underlying the field sites have reported δ^13^C values of 6.7 (± 1.2) ‰ (Bekker et al., 2003). Thus, dissolution of Nash Fork Fm. carbonate with atmospheric CO_2_ would produce a predicted DIC δ^13^C value of −2.5 ‰, again heavier than our snow samples.
3. If the primary source of CO_2_ was from heterotrophic breakdown of organic carbon (C_org_), we can assume CO_2_ δ^13^C values would mirror the Corg source, with examples of soil (C_org_ δ^13^C = −24.1 ‰; Table S1), DOC (C_org_ δ^13^C = −22.2 ‰; Table 1), or bulk contextual plant samples (C_org_ δ^13^C = −25.9 ‰; Table S1). Thus, DIC sourced solely from heterotrophically produced CO_2_ would reflect these values, and DIC sourced from dissolution of Nash Fork carbonate by heterotrophically produced CO_2_ would have estimated δ^13^C values calculated from δ^13^C_HeterotrophicBreakdown_ - δ^13^C_NashCarbonate_.
4. Meltwater discharging from the surface and subsurface of the Round Lake snow field site had a pH of 8.5 and DIC (99.9 μmol/L) with a δ^13^C value of −10.3 ‰, and meltwater discharge from the Jeep Lake snow field site had a pH of 7.5 and DIC (179.4 μmol/L) with a δ^13^C value of −11.0 ‰. This is consistent with dissolution of carbonate (higher pH) with a combination of atmospheric sourced and soil Corg sourced CO_2_ – as the measured DIC δ^13^C values fall between the predicted values of −2.5 and −15.5 ‰.
5. Measured Round Lake snow DIC δ^13^C values were ~ 5 to 10 ‰ lighter than the −9.2 ‰ predicted for purely atmospheric CO_2_ input, which in combination with the lower pH (6.0 to 6.2) this implies an additional input of Corg-derived CO_2_.

Jeep Lake snow DIC returned an extremely negative δ^13^C value (−24.0 ‰), implying DIC input of solely heterotrophically-sourced CO_2_, but this is inconsistent with the relatively alkaline pH (7.7) which necessitates carbonate dissolution buffering. Thus, we suspect that Jeep Lake snow sample DIC δ^13^C values (and also potentially but to a lesser extent Round Lake) were impacted by accelerated production of CO_2_ due to solar heating of collected snow samples required to filter water for geochemical analyses and separate biomass from snow. This impact was noted in two higher biomass snow samples from the PNW – causing DIC δ^13^C values to be 5.4 to 8.3 ‰ more negative than predicted. This highlights a need for caution when collecting geochemical data from melted snow – with care needed to minimize heterotrophic input – which might be remedied through collection of replicates of snow from below the active snow algae surface for pH and DIC analyses to supplement samples of active snow algae snow.

Additional abiotic factors might impact snow algae composition. In general, Round Lake samples had higher concentrations of iron while Jeep Lake samples (except the for lake) had more phosphorus, manganese, calcium, sulfate, and dissolved inorganic carbon (Table 1). pH and geochemistry are strong drivers of community composition in other environments including hot springs (e.g., Brown and Fritz, 2019; Hamilton et al., 2019;), lakes (e.g., Schweitzer-Natan et al., 2019), and soils (e.g., Bahadori et al., 2021). Time of year could also play a key role in controlling snow algae composition as has been observed in other locations (e.g., Hotaling et al., 2022; Winkel et al., 2022). Nutrient concentration, which can also vary seasonally (Winkel et al., 2022), and other factors including precipitation and initial snow depth are likely controls on snow algae composition and abundance throughout the course of the melt season. Even though our samples were collected at similar times (August in 2019 and 2021), we cannot discount seasonal variation as a factor leading to the differences we observe in snow algae communities on yellow to orange snow collected above Round Lake and pink snow collected from above Jeep Lake.

### Microcosms

While nutrients in snow can vary seasonally (Winkel et al., 2022), snow is generally low in aqueous nutrients including fixed nitrogen, phosphorous, and iron (Hamilton and Havig, 2017; Havig and Hamilton, 2019). Round and Jeep Lake snowfields were low in dissolved fixed nitrogen — NH_4_(T) and NO_3_- were below detection limits (NH_4_(T) = 1.4 μM, NO_3_- = ~ 7 nM, respectively)(Table 1). We tested our nutrient limitation hypothesis using a microcosm approach where we added either solely NaH^13^CO_3_ or NaH^13^CO_3_ plus either fixed nitrogen, phosphorus or iron and measured the amount of excess ^13^C taken up in biomass following incubation for two hours. The addition of nutrients in the form of NH_4_+, NO_3_-, PO_4_^3-^ or Fe did not stimulate primary productivity (rates were the same or lower than our “light” control)(Figure 3). These data suggest Jeep and Round Lake snow algae communities were not limited in fixed nitrogen, phosphorus, or iron. While fixed nitrogen (NO_3_- and NH_4_(T)) was at or below detection limits in all samples measured, our samples contained measurable amounts of total dissolved Fe (474 and 1362 nM in Round Lake snow and 390 nM in Jeep Lake snow) and total dissolved P (~0.5 μM in Round Lake snow and 3.3 μM in Jeep Lake snow)(Table 1). We cannot rule out the possibility that snow algae communities are co-limited by multiple nutrients.

**Figure 3.**
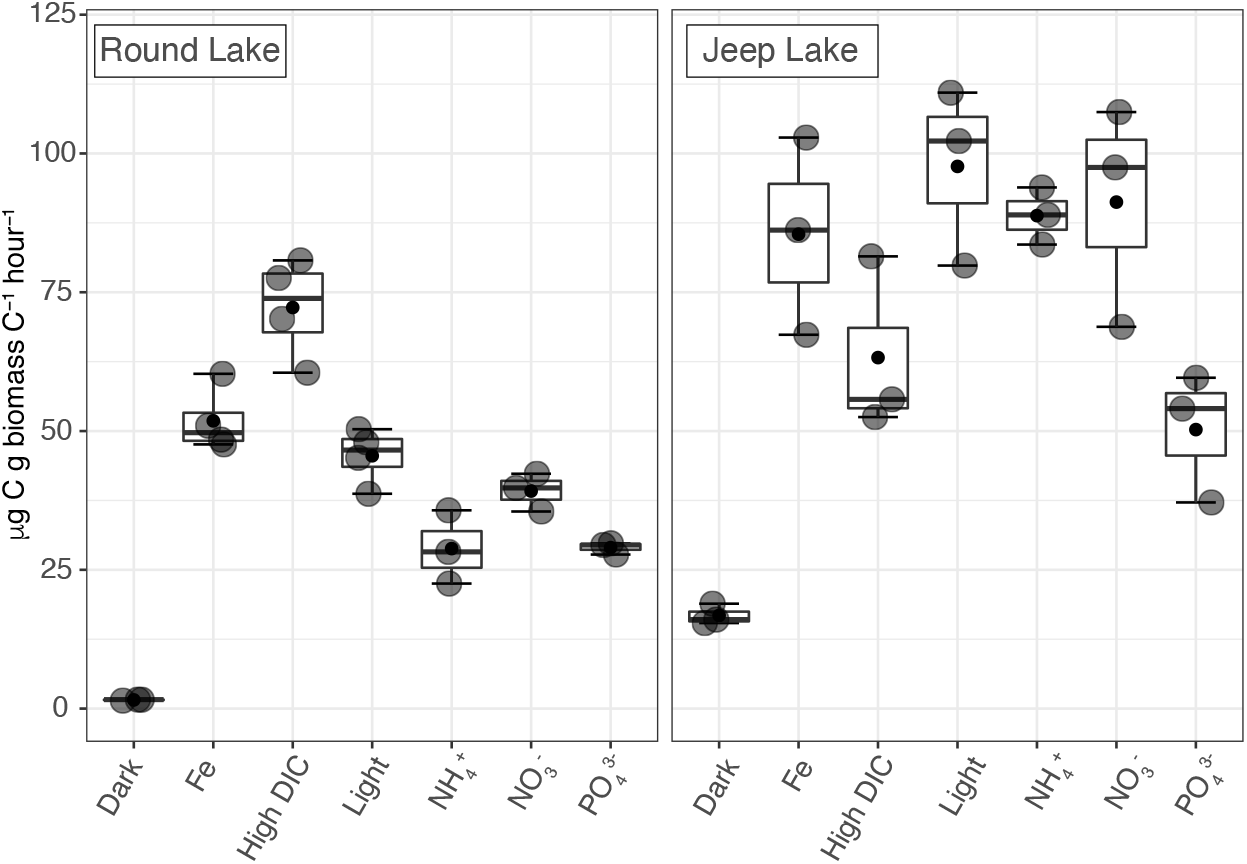
Rates of carbon assimilation in microcosm assays. The horizontal line in each box indicates the median and closed circles represent the mean (*n* = 3 or 4 for each treatment).

In our previous studies of supraglacial snow algae in the PNW, we observed stimulation of snow algae primary productivity upon the addition of inorganic carbon (in the form of NaHCO_3_)(Hamilton and Havig 2020). The concentration of dissolved inorganic carbon in PNW snow and ice are low (≤ 55 μM) suggesting inorganic carbon limits snow algae primary productivity (Hamilton and Havig 2017; 2020; Havig and Hamilton 2019). Primary productivity in Round Lake snow was stimulated by the addition of DIC concentrations – with ‘high DIC’ addition exhibiting the highest productivity increase – while primary productivity Jeep Lake snow microcosms did not have the same response (Figure 4). This begs the question — what is the impact of inorganic carbon source for Round and Jeep Lake snow algae.

**Figure 4.**
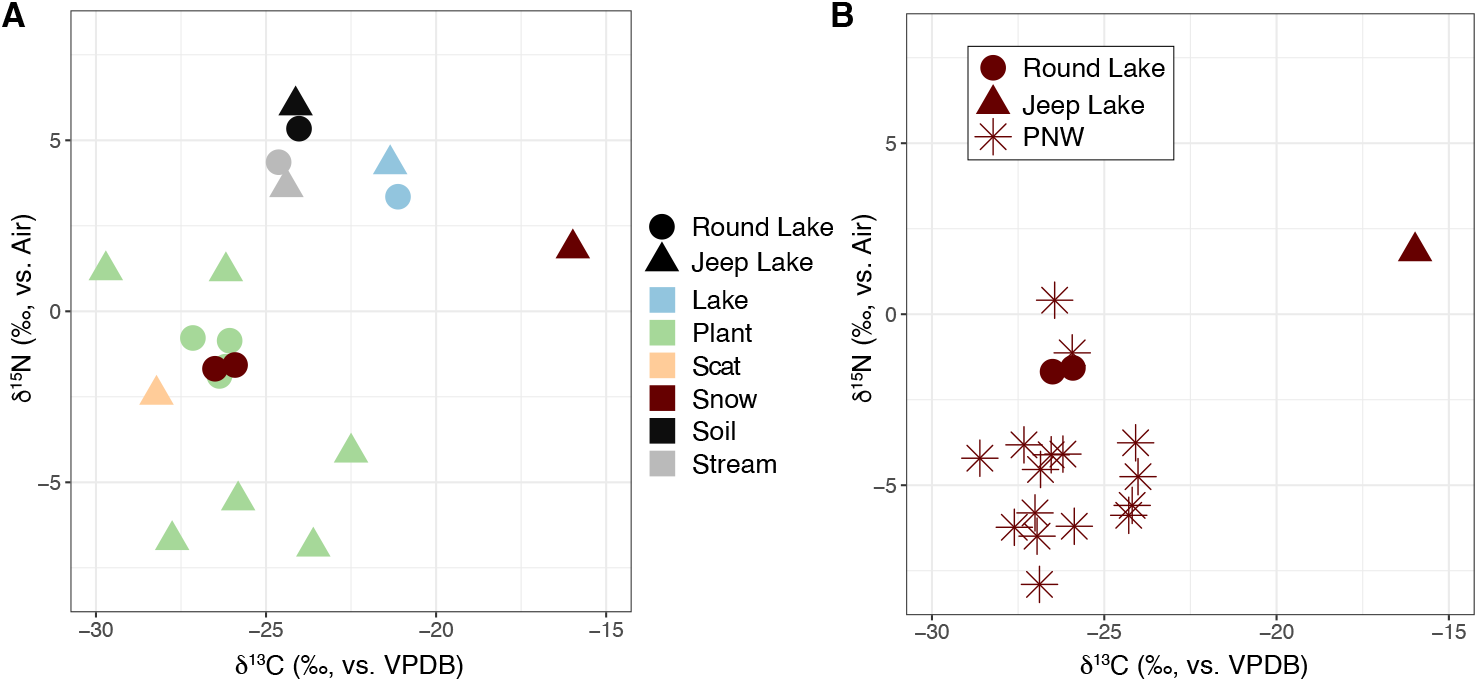
Biomass C and N stable isotope analyses. Values are provided in Tables 2; S1; and S2.

**Figure 5.**
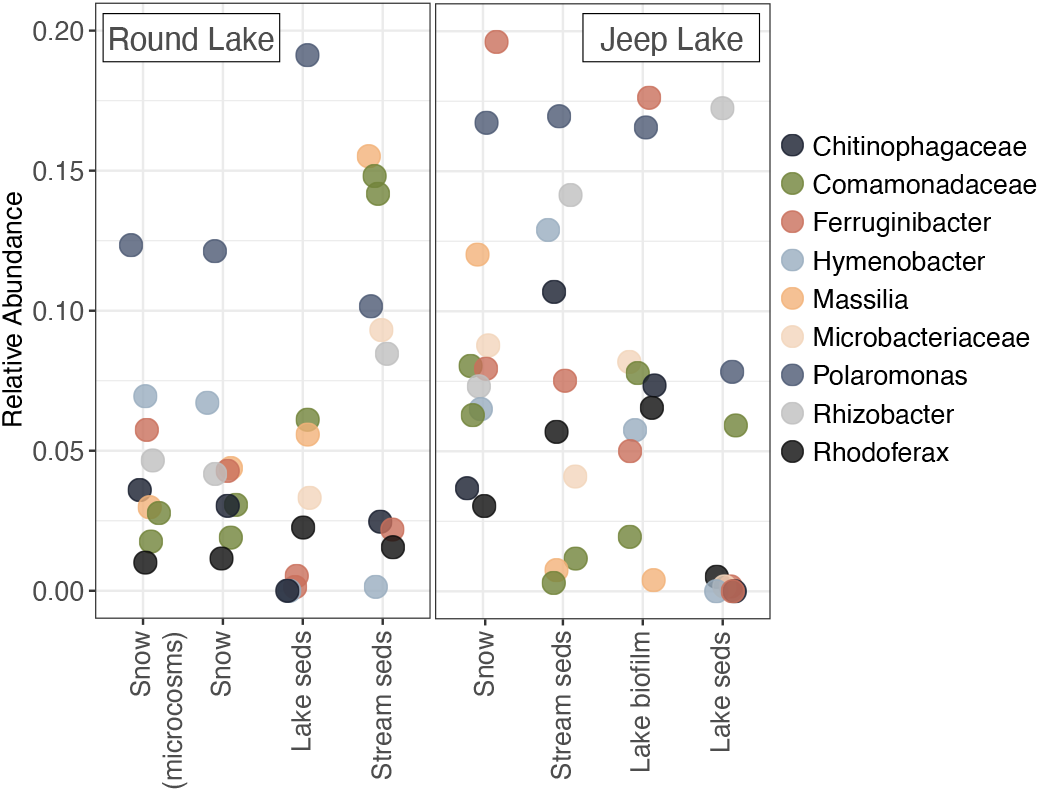
Relative abundance of bacterial small subunit rRNA (16S rRNA) OTU transcripts recovered from snow, lake and stream sediments (seds) and lake biofilms. OTUs for each library were binned at the family level. The Round Lake sample that corresponds to the site where microcosms were performed is indicated.

δ^13^C values of Round Lake snow algae biomass were −26.5‰ and −25.1‰ (Figure 4A; Table 2), consistent with previous values (Hamilton and Havig, 2017, Havig and Hamilton, 2019). In contrast, Jeep Lake snow algae biomass had a δ^13^C of −16.0 ‰ which is heavier than typically expected for algal carbon fixation. The difference between snow algae biomass δ^13^C values at the Round Lake (−25.9 and −26.5 ‰) versus Jeep Lake (−16.0 ‰) sample sites highlights the impact of DIC source for δ^13^C signal and would leave one to predict a difference in response to DIC addition experiments between the two sites. Fractionation factors (Δ^13^C = δ^13^C biomass - δ^13^C DIC) calculated from Round Lake samples (assuming the primary CO_2_ source is atmospheric, with a value of −9.2 ‰) give a Δ^13^C value of 17 (16.7 to 17.3) ‰, which align with PNW results (n = 11) suggesting snow algae Δ^13^C is typically 17.8 (± 1.5) ‰ and is consistent with fractionation factors predicted for the pentose phosphate cycle (Havig et al., 2017; Garcia et al., 2021). Based on these observations, we would predict a DIC source with a δ^13^C value of ~ 1.0 ‰ for the snow algae biomass (δ^13^C of −16.0 ‰) sampled at the Jeep Lake site. This value best aligns with a primary DIC source from atmospheric CO_2_ dissolution of Nash Fork Fm. carbonate (−2.5 ‰). This is consistent with earlier discussion – which suggested the Round Lake snow fields at the time of sampling had atmospheric CO_2_ as a primary source for DIC (low DIC concentration, slightly acidic pH) and were thus potentially carbon limited while Jeep Lake snow fields had alkaline pH values and higher DIC concentrations consistent with dissolution of locally-derived carbonate minerals driven primarily by atmospheric CO_2_ (but also with a heterotrophically-sourced CO_2_ component). This result directly addresses the core hypothesis of this work – testing if snow fields with carbonate-bearing bedrock and soil are alleviated from the DIC limitation observed for snow algae communities hosted on volcanic terrains devoid of carbonate minerals. Our results suggest that while it is possible for carbon limitation to be alleviated from increased DIC availability due to dissolution of carbonate sourced from local bedrock (as was the case for the Jeep Lake snow algae), it is not necessarily a ubiquitous or universal effect (as shown by the increased primary productivity in response to DIC amendment for Round Lake snow algae).

**Table 2.** Total carbon and nitrogen and stable isotope analysis results for Medicine Bow snow, lake, and stream biomass.

| Sample Type | Total C<br>% | $\delta^{13}\text{C}$<br>‰ | Total N<br>% | $\delta^{15}\text{N}$<br>‰ |
| --- | --- | --- | --- | --- |
| Round Lake |  |  |  |  |
| snow algae | 9.03 | -26.50 | 0.51 | -1.68 |
| snow algae | 8.21 | -25.91 | 0.44 | -1.57 |
| stream sediments | 0.94 | -24.63 | 0.09 | 4.36 |
| lake sediments | 0.84 | -21.12 | 0.07 | 3.35 |
| Jeep Lake |  |  |  |  |
| snow algae | 25.00 | -15.98 | 1.68 | 1.84 |
| stream sediments | 2.85 | -24.40 | 0.17 | 3.63 |
| lake sediments | 3.72 | -21.35 | 0.30 | 4.30 |

### Bulk snow algae community biomass nitrogen source

The N content and δ^15^N values of snow algae biomass at Round Lake was 0.44 to 0.51 % and −1.6‰ to – 1.7‰ and the Jeep Lake snow algae was 1.68 % and 1.8 ‰, respectfully (Figure 4A; Table 2). These values overlap with δ^15^N values expected for nitrogen fixed from atmospheric N_2_ (with δ^15^N value of 0‰) via biological N_2_-fixation, which results in δ^15^N values of ~ +2 to −2‰. Jeep Lake snow algae biomass fell at the higher end of the range while Round Lake snow algae fell at the lower end.

Soil δ^15^N values were 5.3 to 6.0 ‰, consistent with enrichment in ^15^N from breakdown of organic matter and preferential loss of ^14^N via nitrogen cycling and denitrification processes. Stream and lake sediment δ^15^N values (3.4 to 4.4 ‰) also reflect similar processes impacting the N isotope signal. Plant δ^15^N values ranged from 1.2 to −6.9, consistent with fixed nitrogen from multiple sources. Plant δ^15^N values from the PNW ranged from 1.8 to −8.9 ‰ (Figure 4B), with more positive values attributed to uptake from soil, values closer to a value of 0 ‰ attributed to a nitrogen fixation signal, and values below – 2 ‰ attributed to uptake of fixed nitrogen from atmospheric deposition (Havig and Hamilton, 2019). Much of the atmospheric deposition-sourced fixed N was attributed to ^14^N-enriched NH3 volatilized from dairy and feed lot sources located upwind from the snow algae sites (Macko and Ostrom, 1994; Hristrov et al., 2009; Deng et al., 2018), as well as potential marine sources (i.e., the Pacific Ocean)(Jickells et al., 2008). Plant δ^15^N values from the Snowy Mountain sites follow similar trends, suggesting similar nitrogen input assimilation patterns. The accumulation of ^14^N-enriched NH_4_^+^ with winter and spring snow accumulation is consistent with previous work (e.g., Russell et al., 1998), and fixed nitrogen δ^15^N values of −2.0 to −5.5 ‰ for precipitation in the Rockies have been reported (Nanus et al., 2008). Snow algae δ^15^N values of - 1.6 to −1.7 ‰ at Round Lake are consistent with a lower soil input component and higher precipitation source input, while δ^15^N values of 1.8 ‰ for Jeep Lake are consistent with a greater soil input component, both of which align with respective C-isotope and uptake results. However, the snow algae δ^15^N values also overlap with the expected range for fixed N sourced from nitrogen fixation, suggesting there might be a contribution from N-fixing microbial community members.

Algae do not fix nitrogen but there are examples of microbes in the 16S rRNA data that may fix nitrogen. For example, species in the *Polaramonas* are known to fix nitrogen (Hanson et al., 2012). OTUs affiliated with *Polaromonas* were abundant in Round Lake and Jeep Lake snow, with *Polaromonas* accounting for 14-15 % of recovered 16S rRNA OTUs in Round Lake snow algae samples and 18 % for Jeep Lake. *Polaromonas* are commonly observed in snow and glacier ecosystems (Mattes et al., 2008; Darcy et al., 2011; Michaud et al., 2012; Hamilton and Havig, 2017; Havig and Hamilton, 2019) including oligotrophic soils in glacier forefields (Nash et al., 2018). The presence of *Polaromonas* in glacier fore field sediments has been linked to accelerated weathering through granite dissolution, resulting in the release of essential trace elements including Mn and Fe (Frey et al., 2010). δ^15^N values of plant material from the surrounding landscape is also consistent with nitrogen from nitrogen fixation while soil δ^15^N was heavier (~ +4 ‰). Thus, plant material delivered to the snow surface may also be a source of fixed nitrogen to snow algae communities — OTUs affiliated with *Ferruginibacter,* which are known to degrade complex carbon (del Rio et al., 2010), were abundant in Jeep Lake snow.

## CONCLUSIONS

Community composition was distinct between two snowfields in the Snowy Range despite being separated by ~500 m. Even with increased nitrogen and phosphorus deposition, ammonia and nitrate concentrations were low in Snowy Range snow and snow algae primary productivity was not stimulated by the addition of aqueous ammonia, nitrate, phosphate, or iron. Similar to our observations in the Cascade Range, addition of inorganic carbon stimulated snow algae primary productivity in the snow with lower inorganic carbon concentration (Round Lake) despite the presence of carbonate bedrock, which alleviated inorganic carbon limitation for the Jeep Lake site. Our results suggest snow algae (at least late in the melt season), are not limited by fixed nitrogen, phosphorus or iron even in areas that might receive higher deposition of the essential nutrients and snow algae productivity, and thus abundance, which is related to melt, can be linked to increasing CO_2_. Furthermore, the presence of carbonate bedrock as a potential source for additional inorganic carbon does not necessarily alleviate carbon limitation of snow algae productivity – suggesting inorganic carbon limitation of snow algae blooms may be a nearly universal pattern and supporting the hypothesis that increased atmospheric CO_2_ concentrations may lead to larger and more robust snow algae blooms globally.

## ACKNOWLEDGEMENTS

This work was supported by the University of Minnesota. The authors acknowledge the Minnesota Supercomputing Institute (MSI) at the University of Minnesota for providing resources that contributed to the research results reported within this paper. The authors acknowledge that this work was conducted on the ancestral lands of the hinono’eino’ biito’owu’ (Arapaho), Newe Sogobia (Eastern Shoshone), and Tséstho’e (Cheyenne) (source: https://native-land.ca/) and support all efforts to give these tribes a voice and power to guide decisions in management practices for the Snowy Mountain Range.

## DATA AVAILABILITY

Raw reads are available on NCBI under accession numbers PRJNA936880.

## AUTHOR CONTRIBUTIONS

T.L.H. and J.R.H. designed the study, collected field samples, and perform field microcosms. T.L.H. analyzed the data and wrote the paper with input from J.R.H.

## MATERIALS & CORRESPONDENCE

Correspondence and requests for materials should be addressed to T.L.H.: Trinity L. Hamilton. Department of Plant and Microbial Biology, University of Minnesota, St. Paul, USA, 55108. Phone: +16126256372.

