## Supplemental Tables for "Carbonate bedrock may not alleviate DIC limitation of snow algae — a test of hypothesis in the Medicine Bow Mountains, WY, USA"

**Table S1**. Total carbon and nitrogen and stable isotope analysis results for context biomass samples. Values in grey are near the limits of calibration curves, and should be taken as approximations.

| Sample Type | Total C % | $\delta^{13}C$ ‰ | Total N % | $\delta^{15}N$ ‰ |
|---|---|---|---|---|
| Round Lake | | | | |
| soil | 1.60 | -24.03 | *0.11* | *5.34* |
| grass | 42.10 | -27.15 | 2.96 | -0.78 |
| clover | 44.80 | -26.19 | 3.10 | -1.62 |
| spruce needles (living) | 48.44 | -26.37 | *1.20* | *-1.89* |
| dead needles | 47.32 | -26.07 | *1.23* | *-0.86* |
| Jeep Lake | | | | |
| soil | 1.62 | -24.13 | 0.12 | 6.02 |
| grass | 8.67 | -22.50 | 0.44 | -4.13 |
| sedge | 42.83 | -26.18 | 2.70 | 1.17 |
| elephant flower leaves | 47.38 | -27.76 | 2.38 | -6.69 |
| spruce needles (living) | 47.91 | -25.82 | *1.16* | *-5.53* |
| moose poop | 43.58 | -28.22 | 2.05 | -2.44 |
| moss | 20.66 | -23.61 | 2.32 | -6.88 |
| huckleberry bush leaves | 47.46 | -29.71 | 2.03 | 1.20 |

**Table S2**. Total carbon and nitrogen and stable isotope analysis results for snow algae samples from the PNW. Data from Hamilton and Havig 2017; 2020; Havig and Hamilton, 2019.

| Sample Type | Glacier | Total C % | $\delta^{13}C$ ‰ | Total N % | $\delta^{15}N$ ‰ |
|---|---|---|---|---|---|
| snow algae (red) | Gotchen | 5.94 | -24.19 | 0.328 | -5.59 |
| snow algae (orange) | Gotchen | 6.81 | -28.61 | 0.512 | -4.21 |
| snow algae (red) | Gotchen | 1.90 | -26.44 | 0.162 | 0.41 |
| snow algae (red) | Gotchen | 0.56 | -25.93 | 0.041 | -1.13 |
| snow algae (red) | Eliot | 0.35 | -26.95 | 0.024 | -6.49 |
| snow algae (red) | Eliot | 0.64 | -27.01 | 0.068 | -5.81 |
| snow algae (red) | Eliot | 0.61 | -27.61 | 0.062 | -6.23 |
| snow algae (red) | Diller | 0.08 | -25.92 | 0.007 | bdl |
| snow algae (orange) | Diller | 0.63 | -27.33 | 0.049 | -3.82 |
| snow algae (red) | Collier | 0.48 | -24.02 | 0.025 | -4.75 |
| snow algae (red) | Palmer | 4.01 | -24.29 | 0.209 | -5.88 |
| snow algae (red) | Palmer | 3.29 | -25.87 | 0.156 | -6.20 |
| snow algae (red) | Gotchen | 6.14 | -24.09 | 0.410 | -3.76 |
| snow algae (red) | Gotchen | 29.66 | -26.20 | 1.475 | -4.09 |
| snow algae (red) | Eliot | 0.98 | -26.85 | 0.093 | -4.54 |
| snow algae (red) | Eliot | 0.44 | -26.88 | 0.054 | -7.90 |
| snow algae (red) | Collier | 0.97 | -26.54 | 0.064 | -4.11 |
